# Biotinylation-dependent near-neighbor analysis for identification of septation regulators in *Aspergillus fumigatus*

**DOI:** 10.64898/2026.09.17.752449

**Authors:** Harrison Thorn, Adela Martin-Vicente, Uxue Perez-Cuesta, Jinhong Xie, Devi Bale, Asmita Nandi, Jarrod R. Fortwendel

## Abstract

Recent work has established septation as a critical process for virulence in *A. fumigatus,* as strains lacking septa are avirulent. While our work has focused on the well-studied Septation Initiation Network (SIN), understanding of septation machinery upstream and downstream of SIN remains limited in filamentous fungi. Proximity labeling techniques are powerful tools to study pathway interactions, especially those that are transient. Recent advances have seen TurboID used to study pathways and processes important for fungal pathogenesis. In this study, we adapt TurboID for use in *Aspergillus fumigatus* and use overlapping datasets of two SIN components, the terminal NDR kinase SidB and its binding partner MobA, to identify putative septation effectors. Many known septation effectors, including septins and cell wall synthesis proteins, were enriched in our datasets, validating the approach. Phenotypic characterization of hits identified by LC-MS/MS revealed several previously uncharacterized gene products involved in septum formation and cell wall stress tolerance. These included the PP2A regulatory subunit PabA and IQGAP SepG, which have been shown to regulate septation in *Aspergillus nidulans*. Thus, TurboID is a useful tool to study fungal signaling and physiology in *A. fumigatus* and may be used to improve understanding of pathogenic processes.

## Introduction

*Aspergillus fumigatus* is the major filamentous fungal pathogen causing invasive fungal infections [1]. Our group and others have recently observed loss of septation correlates with a decrease in virulence and pulmonary tissue invasion in mouse models of Invasive Pulmonary Aspergillosis (IPA) and confers fungicidal activity to the echinocandin class of antifungal drugs [2–4]. Therefore, septation inhibition may represent a novel therapeutic strategy to both blunt *A. fumigatus* virulence and improve antifungal killing of the echinocandins, which are used only as salvage therapy to treat invasive *Aspergillus* infections in part due to their fungistatic activity [5, 6]. Septation in *A. fumigatus* and related fungi is orchestrated by a conserved kinase cascade called the Septation Initiation Network (SIN). Many SIN components are essential for septation, necessitating further study of this pathway [2, 3, 7]. Much of what is known about SIN signaling has been elucidated in the model fission yeast *Schizosaccharomyces pombe* (reviewed in [7]). Beginning in anaphase, the GTPase Spg1p becomes activated at the spindle pole body (SPB), driving stepwise signaling through three kinases, Cdc7p, Sid1p and Sid2p [8–10]. Sid1p and Sid2p are further activated via binding to required cofactors Cdc14p and Mob1p, respectively [11, 12]. Following Sid2p activation, it travels from the SPB to the septation site alongside its required coactivator Mob1p [13]. Orthologs of these proteins in *A. fumigatus* are known as SpgA (Spg1p), SepH (Cdc7p), SepL (Sid1p), SepM (Cdc14p), SidB (Sid2p) and MobA (Mob1p), and have been characterized by our group. As in *S. pombe*, the three SIN kinases and MobA are essential for septation in *A. fumigatus*. While not strictly required, SepM is a critical septation mediator, whereas SpgA was dispensable for septation in *A. fumigatus* [2, 3]. Our previous work indicated that gene deletion of SIN kinases or coactivators conferred enhanced killing activity to the echinocandin antifungals, compared to their usual fungistatic activity while simultaneously reducing virulence in animal models of IPA. [2, 3, 6]. We reasoned that due to its localization at both the SPB and the septation site, the SidB/MobA module likely initiates key events early in septum formation (i.e. cytoskeletal organization and septal wall synthesis) [12, 14]. As these signaling steps may be critical for virulence and echinocandin sensitivity, we aimed to identify near neighbors of the SidB/MobA module, which remain poorly understood in filamentous fungi.

As a kinase, interactions between SidB and its upstream activators and downstream substrates may be transient, precluding the usage of traditional immunoprecipitation to pursue this goal. Therefore, we aimed to adapt a proximity-dependent labeling (PDL) tool to identify protein-protein interactions that putatively regulate SidB/MobA signaling. PDL usage is increasing in fungi, typically harnessing the strong biotin-streptavidin interaction to study protein-protein interactions in a cell [15–17]. Biotin-based PDL involves fusing a protein of interest to a promiscuous biotin ligase derived from *Escherichia coli* BirA. BirA was modified to BioID, with improved efficiency, and later to TurboID, which acts much faster than BioID [18]. The bait-ligase chimera catalyzes the conversion of biotin to the reactive biotinoyl-5’-AMP anhydride, which reacts with free lysine residues on neighboring proteins, generating a covalent biotin signature [19]. By creating a covalent label, near neighbors of the bait remain identifiable even in the absence of a direct interaction, so this is a favorable system for studying short-lived interactions on the time course of a phosphorylation event [18]. After the labeling period, biotinylated prey is isolated from crude lysates via pulldown with streptavidin-conjugated magnetic beads. Biotinylated proteins are then eluted and analyzed by liquid chromatography followed by tandem mass spectroscopy (LC-MS/MS).

Here, we establish a TurboID system for proximity-based labeling and detection in the opportunistic fungal pathogen *A. fumigatus*. Using TurboID-conjugated SidB/MobA, we successfully identified known septation regulators, validating the system. Downstream analysis of previously uncharacterized proteins revealed conserved septation effectors as putative SidB/MobA neighbors. Several of these displayed increased echinocandin sensitivity consistent with septation deficiency, potentially guiding therapeutic development. This work emphasizes the utility of PDL to study disease-associated traits of human fungal pathogens.

## Results

### Establishment of TurboID in *A. fumigatus*

To produce *A. fumigatus* strains that could be used for biotin-based PDL approaches, we first utilized CRISPR/Cas9 gene editing to generate in-frame insertions of the TurboID coding sequence at the 3’ end of the *sidB* and *mobA* genes (Figure 1A). As our mutational approach targeted integration at the native loci for each gene, expression of the resulting SidB-TurboID and MobA-TurboID chimeric proteins is driven by the native promoter. These genetic manipulations also included the addition of an 18-base pair linker sequence upstream of TurboID to minimize steric hindrance of the resulting chimeric protein. For controls, we predicted that two separate strains should be highly informative. First, we constructed a strain expressing the TurboID gene from a known safe-haven locus under control of the constitutive pOtef promoter [20, 21] (Figure 1B). This strain, expressing free TurboID, would serve as the analytical background to quantify significantly enriched interactions for the SidB-TurboID and MobA-TurboID strains in downstream proteomics. To ensure our approach could identify pathway-specific interactions, a second control strain was constructed in which MobB, an NDR kinase co-activator of the distinct RAM signaling pathway, was also fused with TurboID as previously described for the SIN components [22]. We reasoned that the MobA-TurboID (SIN pathway) and MobB-TurboID (RAM pathway) chimeras should largely biotinylate different subsets of proteins and therefore identify distinct interactomes.

**Figure 1:**
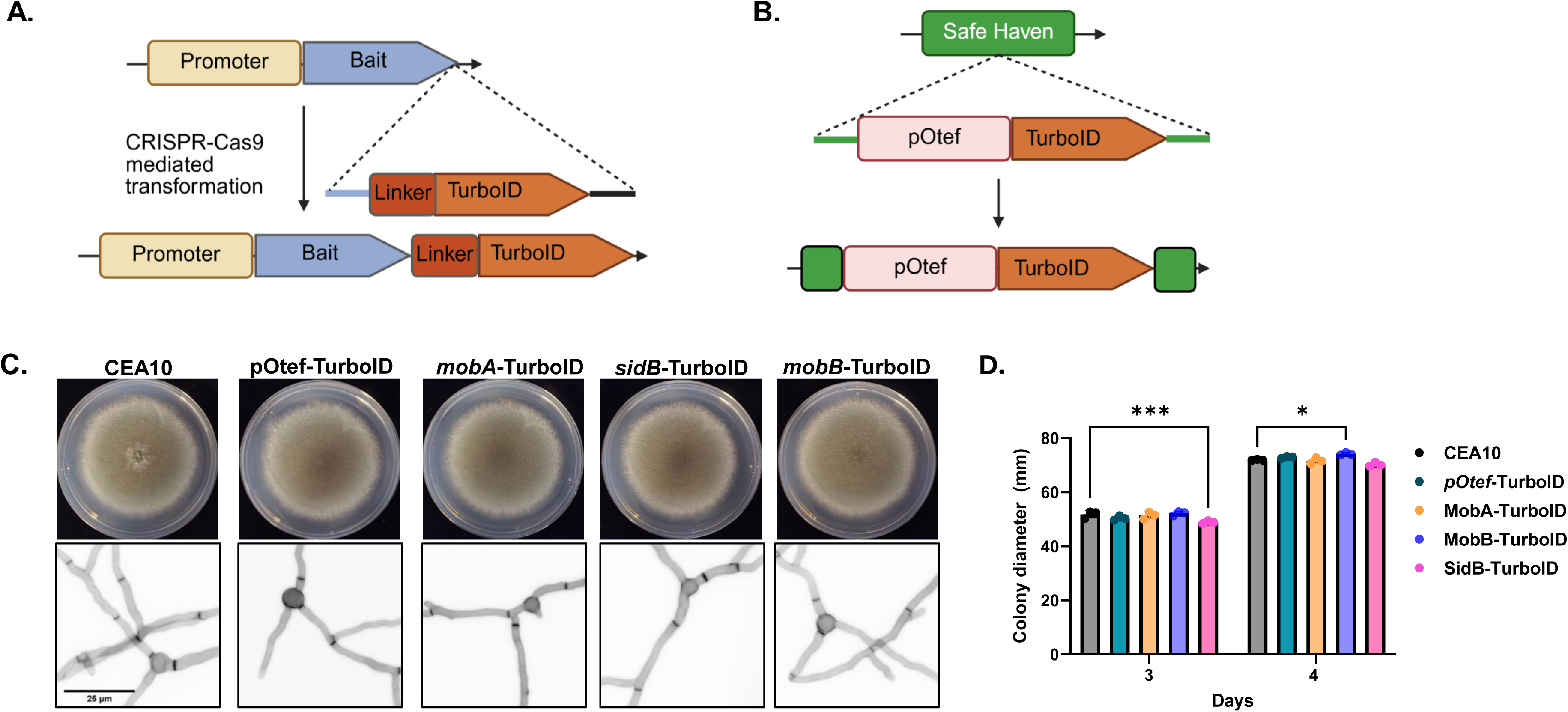
Construction of TurboID chimeras in *A. fumigatus*. **A)** Schematic describing construction of strains expressing chimeric proteins containing a C-terminal TurboID tag. **B)** Schematic describing construction of pOtef-TurboID by insertion of the repair template at a previously annotated “Safe Haven” region in the *A. fumigatus* genome. **C)** 10^4^ conidia were spot inoculated onto solid GMM agar (top) or into GMM broth in a glass bottomed 24-well plate (bottom) to visualize colony and septal morphologies, respectively. For colony morphology, plates were incubated at 37°C for 4 days before imaging. To visualize septa, cultures were incubated at 37°C for 16 hrs prior to calcofluor white staining and imaging on an inverted fluorescent microscope using a DAPI filter. All images are representative of 3 biological replicates. **D)** Diameter of colonies grown as in **(C)** were measured after 3 and 4 days incubation. Average colony diameters (n = 3) were compared to that of CEA10 by two-way ANOVA with Dunnett’s multiple comparison test. ***: p=0.001. *: p<0.05.

To verify that TurboID fusion did not disrupt bait protein function, we performed a radial growth assay to evaluate colony formation and calcofluor white (CFW) staining of mature hyphae to test for proper septation. We previously reported that genetic loss of SidB or MobA causes formation of white colonies and complete loss of hyphal setation [2, 3]. In contrast, the colony morphology, color and diameter of the SidB-and MobA-TurboID strains were similar to the parental background stain (CEA10), suggesting that these fusion events were not detrimental to overall growth or to bait protein function (Figure 1C and 1D). As MobB is a conserved co-activator required for activity of the NDR kinase CotA, and we have previously shown that CotA is an essential kinase in *A. fumigatus*, we predicted that the *mobB* gene would also be essential [22, 23]. In agreement with this prediction, we were unable to generate a *mobB* deletion strain after multiple attempts. However, placing *mobB* expression under the control of a doxycycline regulatable promoter revealed that *mobB* repression resulted in heavily stunted growth supporting its likely essential nature (Supplemental Figure 1). Again, in contrast, the MobB-TurboID strain grew similar to the parental control suggesting that the fusion did not negatively affect MobB function (Figure 1C and 1D). All TurboID-fused bait strains also formed regular septa when compared to the wild type (Figure 1C). Importantly, we also found that the pOtef-TurboID strain grew similar to the parental control suggesting that expression of free TurboID under the conditions tested here did not disrupt growth (Figure 1C and 1D). As all TurboID-expressing strains appeared wild-type-like in these assays, we concluded that TurboID fusion did not disrupt SidB, MobA, or MobB protein function.

To validate that TurboID-based biotin labeling of proteins would be functional in *A. fumigatus*, we next performed a western blot probing crude lysates from the parental background, CEA10, the pOtef-TurboID control, and the SidB-TurboID strains using streptavidin conjugated with horseradish peroxidase (Streptavidin-HRP) to detect biotinylated proteins. Whereas multiple bands were detected in the CEA10 sample, likely representing endogenously biotinylated proteins, blotting of the pOtef-TurboID lysate revealed an increase in biotinylated protein compared to CEA10 (Figure 2A). Interestingly, the SidB-TurboID lysate displayed a level of biotinylation similar to CEA10 (Figure 2A). We reasoned that the differences in detectable biotinylated proteins may be due to relative differences in promoter strengths between the strong pOtef promoter and the native SidB promoter. RT-qPCR performed on RNA extracted from the pOtef-TurboID and SidB-TurboID strains confirmed that the pOtef promoter drives significantly higher expression of TurboID than the native *sidB* promoter (Supplemental Figure 2).

**Figure 2:**
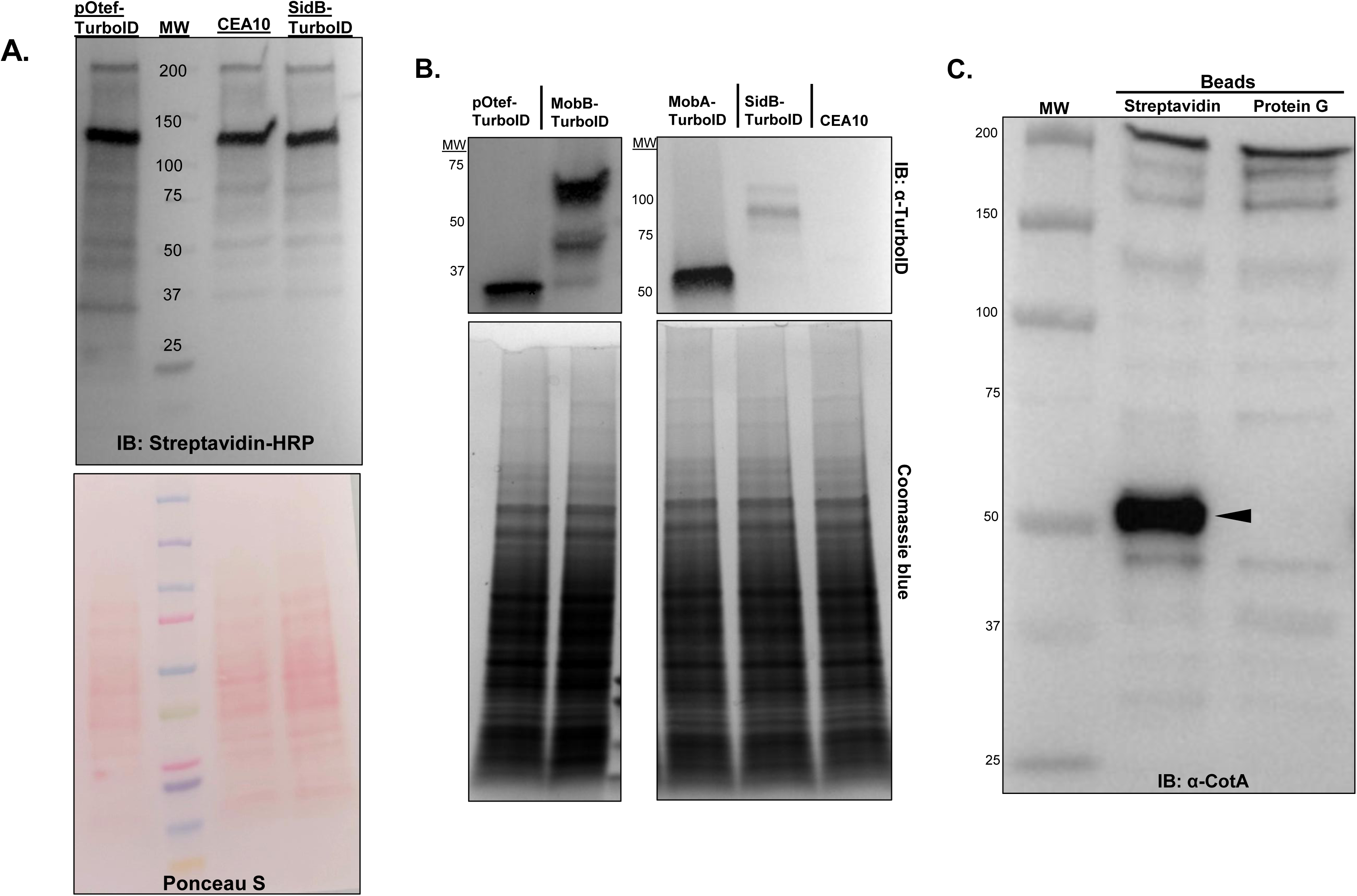
Establishment of TurboID-based proximity labeling in *A. fumigatus*. **A)** 20 μg of total protein lysate was separated on a 4-15% gradient polyacrylamide gel and transferred to a PVDF membrane. The membrane was stained with Ponceau S (bottom) to verify efficiency of transfer. Subsequently, the membrane was blocked with 1% casein in TBS overnight, then probed with streptavidin-HRP (top). **B)** 5mg total protein was incubated with magnetic streptavidin beads for 1 h at room temperature. Proteins were eluted in Laemmli buffer and run on a 4-15% gradient gel, then transferred to a PVDF membrane. The membrane was probed with a TurboID-specific antibody (top). Expected sizes: *pOtef-TurboID*: 35d. MobB-TurboID: 88kd. MobA-TurboID: 67kd. SidB-TurboID: 110kd. **C)** 3mg protein lysate from MobB-TurboID was incubated with magnetic beads coated with streptavidin or protein G for 1 h at room temperature. Proteins were eluted off beads in Laemmli buffer and run on a 4-15% gradient gel, then transferred to a PVDF membrane. The membrane was blotted with a CotA-specific antibody. Only the short isoform was visible, expected size of 50kd.

The fusion of a promiscuous biotin ligase to a bait protein would be expected to result in biotinylation of the bait due to its proximity. Therefore, we next performed precipitations of lysates from each strain using streptavidin beads and utilized these samples for western blot analyses employing an α-TurboID antibody to detect if our TurboID-fused bait proteins were biotinylated. TurboID or TurboID-fused bait proteins were detected in each precipitate, indicating that the chimeras were functional biotin ligases (Figure 2B). To finally validate that our TurboID constructs would biotinylate near neighbors in a predictable bait-specific manner, we took advantage of an α-CotA antibody previously developed by our laboratory to test if TurboID could identify the essential CotA/MobB interaction [22]. Precipitation of crude protein extracts from MobB-TurboID were performed with magnetic beads coated with streptavidin or Protein G (as a control), separated by PAGE, transferred to a membrane, and probed using α-CotA. The CotA protein was clearly visible in streptavidin-based pulldowns of MobB-TurboID lysate but absent in Protein G bead pulldowns (Figure 2C), indicating the ability of TurboID to identify a known interaction. Taken together, our results show that TurboID fusion results in functional chimeras that are able to identify specific interactions, validating the system.

### TurboID analyses detect distinct interactomes for MobA and MobB

To identify proteins proximal to SidB-, MobA-, and MobB-TurboID, crude protein extractions were performed in triplicate after overnight culture of each strain in the presence of 50 μM biotin [24]. pOtef-TurboID was included as a background control. As performed for western blot analyses above, lysates were incubated with streptavidin beads, washed, and precipitated proteins were eluted in the presence of excess biotin. Eluates were processed by LC/MS-MS for protein identification. Principal Component Analysis of all proteins represented by two or more peptides in each sample revealed clearly distinct profiles between each TurboID-fused bait protein and pOtef-TurboID (Figure 3A). Additionally, MobA-TurboID and SidB-TurboID clustered very closely together and were distant from the MobB-TurboID samples, suggesting our chimeric baits performed pathway-specific labeling. To begin decoding pathway-specific TurboID activity, we first defined the experimental interactome of each bait as the collection of protein species in each tagged strain with at least two identified peptides and at least 1.5-fold increased in abundance compared to pOtef-TurboID (p<0.05). This prioritization resulted in interactomes that consisted of 654, 583, and 393 proteins for SidB-TurboID, MobA-TurboID and MobB-TurboID, respectively (Figure 3B –3D). To interrogate these datasets for the ability of TurboID to identify distinct libraries of proteins from bait proteins known to signal in separate pathways, the MobA and MobB experimental interactomes were cross-referenced and found to identify only 118 proteins overlapping in both (Figure 4A). This overlap represents 20.2% and 30% of the MobA and MobB interactomes, respectively. Notably, the MobB binding partner CotA (XP_751641.1) was 99.1-fold overrepresented in MobB-TurboID dataset, while CotA abundance in MobA-TurboID was not significantly different from control (Supplemental Tables 7 and 8). Similarly, the MobA-associated kinase SidB (XP_747793.1) was 57.1-fold overrepresented in MobA-TurboID lysates with no differential abundance in MobB-TurboID pulldowns (Supplemental Tables 7 and 8). Gene ontology (GO) analysis was performed in FungiDB to identify enriched biological processes in MobA-TurboID and MobB-TurboID interactomes (Figure 4B and Figure 4C). GO analysis with MobA-TurboID revealed an enrichment in categories such as anatomical structure development, protein-containing complex assembly, and cytoskeleton organization, consistent with a split localization in complex with SPB scaffolds and the contractile actin ring at the septation site (Figure 4B, Supplemental Table 14) [25]. Consistent with a putative roll for MobB in morphogenesis, categories of anatomical structure development, cell differentiation, and mitochondrial organization were enriched in the MobB-TurboID dataset (Figure 4C, Supplemental Table 15) [22]. This analysis confirmed that a biotin-based near-neighbor approach with MobB-TurboID and MobA-TurboID resulted in the enrichment of proteins in distinct pathways despite their functional similarities, further validating the use of TurboID in *A. fumigatus*.

**Figure 3.**
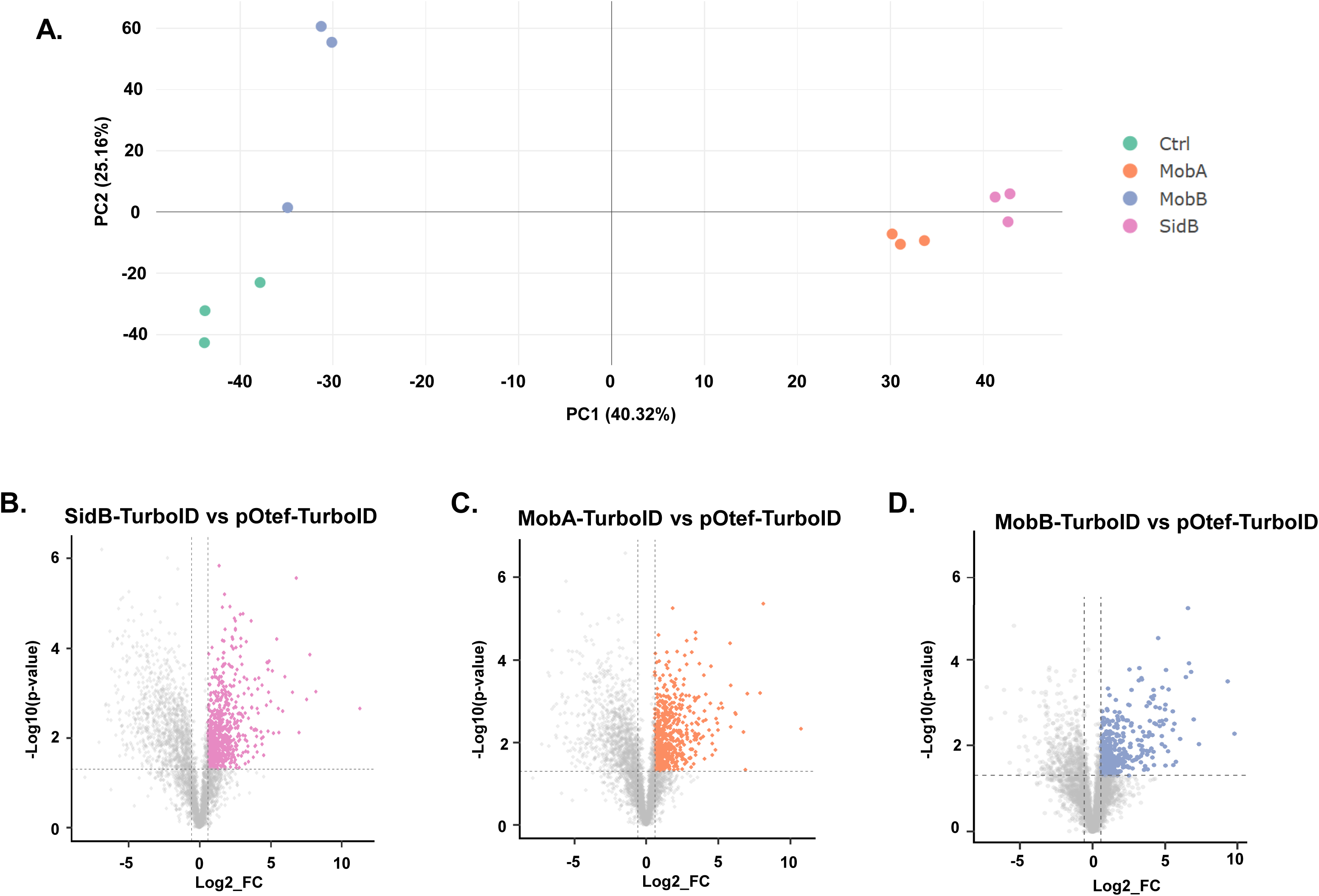
The NDR Kinase regulators, MobB and MobA, are associated with unique interaction networks. **A)** Principle component analysis comparing datasets using the set of TurboID-expressing strains. **B-D)** Volcano plots showing overrepresented proteins in SidB-TurboID, MobA-TurboID, and MobB-TurboID, respectively, each compared with pOtef-TurboID (FC>1.5, p<0.05).

**Figure 4:**
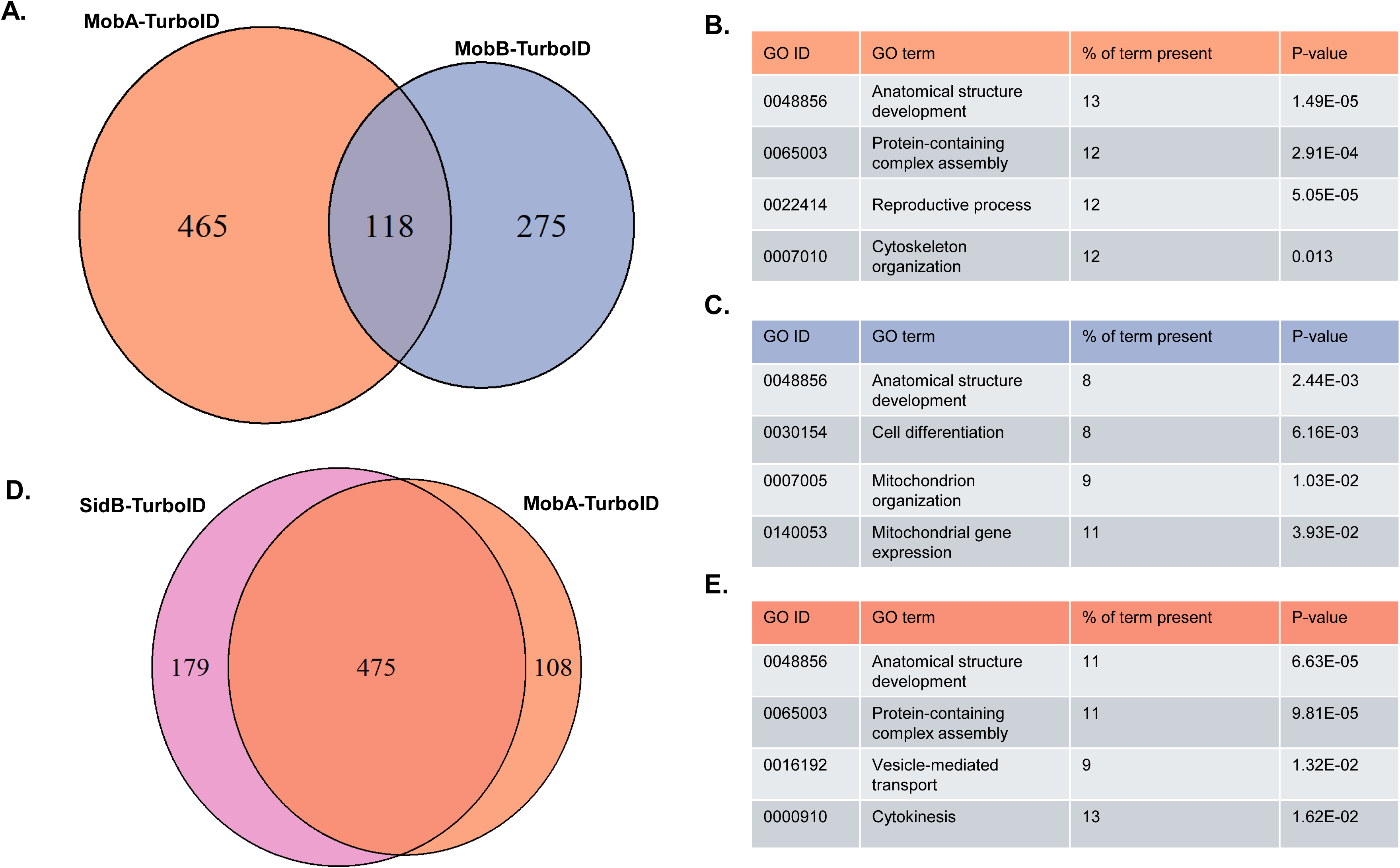
MobA-TurboID and SidB-TurboID enrich a nearly identical proteome. **A)** Venn diagram showing the unique and shared proteins (overlap) identified in MobA-TurboID and MobB-TurboID datasets. **B)** Selected enriched categories determined by GO slim analysis for biological function of proteins associated with proteins enriched in the MobA-TurboID interactome. **C)** Selected enriched categories GO slim analysis for biological function of proteins associated with proteins enriched in the MobB-TurboID interactome. **D)** Venn diagram showing the unique and shared proteins (overlap) identified in MobA-TurboID and SidB-TurboID datasets. **E)** Selected enriched categories GO slim analysis for biological function of proteins associated with proteins enriched in both MobA-and SidB-TurboID interactome.

### Delineation of the SidB/MobA interactome identifies septation regulators

To identify potential regulators and effectors of the SidB/MobA network for further analyses, the MobA-TurboID and SidB-TurboID experimental interactomes were cross-referenced revealing 474 overlapping interactors. This represents 72% and 81.4% of the SidB-TurboID and MobA-TurboID interactomes, respectively (Figure 4D). As SidB was enriched in MobA-TurboID, we identified the reciprocal interaction, with MobA (XP_755571.1) enriched in SidB-TurboID by 6.4-fold, while MobB (XP_752284.1) was 13.4-fold less abundant in SidB-TurboID compared to pOtef-TurboID (Supplemental Table 9). GO analysis of the 474 proteins in both interactomes indicated similar enrichment of biological process categories identified in MobA-TurboID alone. However, inclusion of proteins in both datasets revealed an enrichment in cytokinesis and vesicle-mediated transport consistent with roles in septum formation (Figure 4E and Supplemental Table 17) [26]. Among the proteins represented in these biological processes are multiple known septation regulators in *A. fumigatus*, including the septins AspA (XP_753783.1, 7.5-fold up in SidB, 7.1-fold up in MobA), AspB (XP_748972.1, 3.4-fold up in SidB, 3.2-fold up in MobA), AspC (XP_748035.1, 5.5-fold up in SidB, 5.0-fold up in MobA), and AspD (XP_752252.1, 3.4-fold up in SidB, 3.0-fold up in MobA) [27] (Supplemental Tables 8 and 9). Together, these outcomes suggest a robust experimental SidB/MobA interactome.

We next prioritized uncharacterized proteins in the SidB/MobA interactome for further study to identify components that might be essential for septation. To do so, we performed BLASTp analyses of proteins represented by the highest abundance of identified peptides and lowest p-values. Thirteen proteins with predicted roles in cytoskeletal arrangement and cell wall synthesis/organization were prioritized for gene deletion, as these functions are critical to septum construction [14]. As all were uncharacterized in *A. fumigatus* at the time the work was performed, these thirteen genes were named based on the closest ortholog in *S. cerevisiae* unless an ortholog had been named and characterized in another *Aspergillus* species. Although deletion was successful for all thirteen prioritized genes, the deletion strain of AFUB_020720 (*sepG*) did not produce adequate conidia to perform standard phenotypic characterization experiments. Therefore, we instead inserted the tetracycline-inducible promoter (TetOn) upstream of the predicted *sepG* start codon to generate a strain with conditional gene expression. Growth of this strain under inducing conditions (i.e., with doxycycline) resulted in the return of conidial production (Supplemental Figure 3). Since aseptate strains are often moderately growth restricted and form white colonies, we first performed radial growth assays on minimal media to measure both phenotypes [2, 28] (Figure 5A and 5B). A majority of the mutants displayed statistically significant radial growth defects after 4 days, with Δ*rcoA* (predicted WD-repeat transcriptional repressor) and Δ*pabA* (predicted PP2A regulatory subunit) being the most severely restricted (Figure 5) [29, 30]. These strains, along with Δ*cwf15* (predicted Prp19 spliceosome component) and *sepG^teton^* (conserved IQGAP ortholog) formed colonies with reduced pigmentation, indicative of a reduction in conidiation consistent with a septation defect [31, 32].

**Figure 5:**
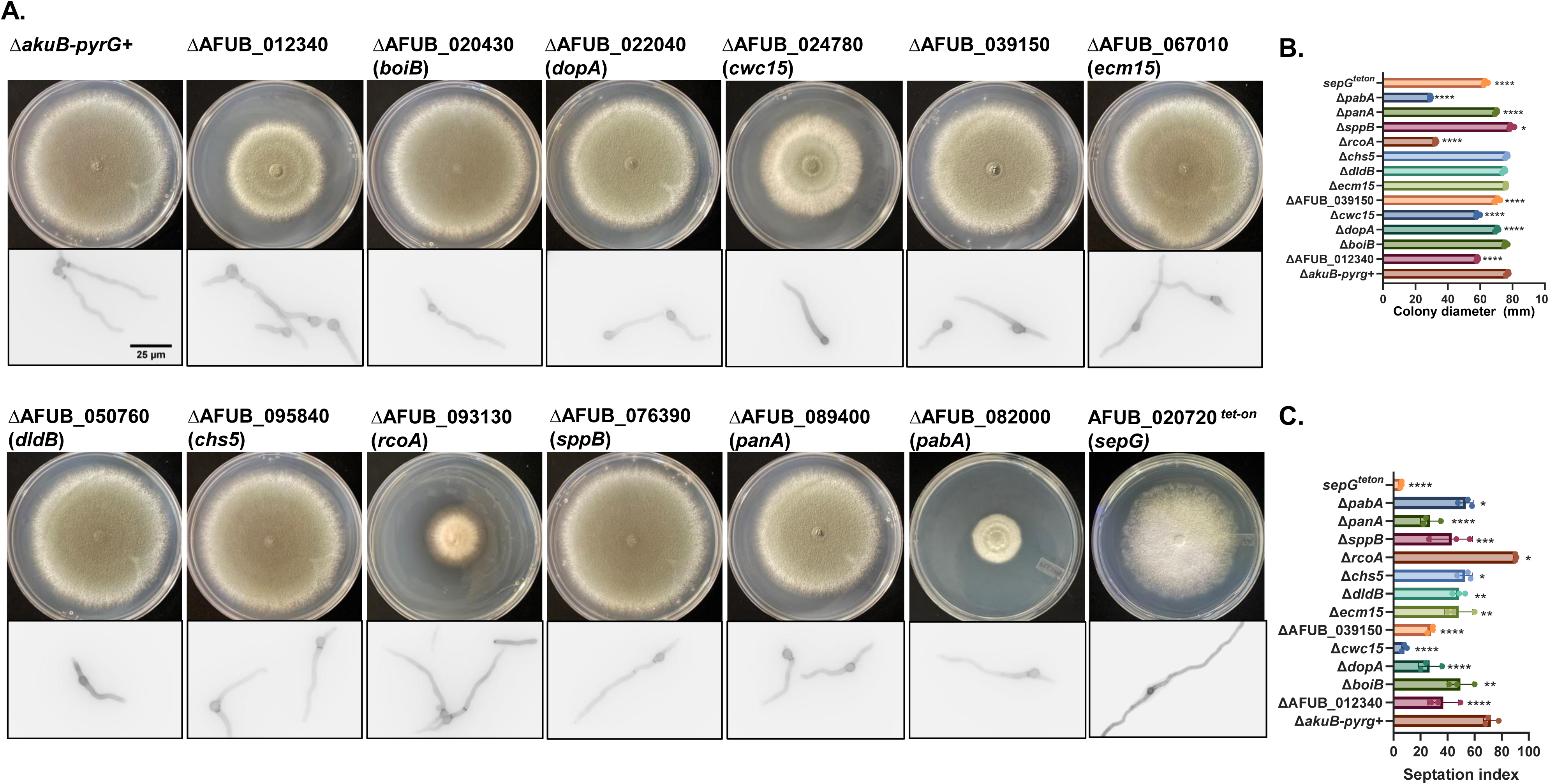
Phenotypic characterization of selected SidB/MobA proximal proteins. A) 10^4^ conidia of the indicated strain were spot inoculated onto solid GMM (top) or into liquid GMM in a glass bottomed 24-well plate (bottom) to visualize colony and septa morphologies, respectively. For colony morphology, plates were incubated at 37°C for 4 d before imaging. To measure septation index, cultures were incubated at 37°C for 10 h prior to CFW staining and imaging on an inverted fluorescent microscope using a DAPI filter. All images are representative of 3 biological replicates. B) Quantification of colony diameter after 4 d. Average diameter was compared to ΔakuB-pyrG+ and analyzed by one-way ANOVA with Dunnett’s multiple comparisons test. *:p<0.05. ****:p<0.0001. C) Quantification of septation index. Average septation index was compared to ΔakuB-pyrG+ and analyzed by one-way ANOVA with Dunnett’s multiple comparisons test. *:p<0.05. **:p<0.01. ***:p<0.001 ****:p<0.0001.

To evaluate if reductions in growth and conidiation were correlated with a loss of septation, we next cultured all strains to an early stage of nascent hyphal formation and stained with calcofluor white to measure the septation index [3]. At this early timepoint, all strains appeared able to form septa but with varying rates of initiation. Although defects in germination rate and hyphal growth in many of these strains could contribute to the variability of septation timing, Δ*cwf15*, *sepG^teton^* (with no doxycycline)*, and* Δ*sppB* had significantly reduced septation indices with only mild to moderate reductions in radial growth rate (Figure 5B and Figure 5C). To test if early septation defects translated into significantly altered septation in mature growth stages, cultures were extended to mature hyphal growth stages and again stained with calcofluor white. With the exception of *sepG^teton^*, all strains exhibited septation albeit at potentially different regularities (Supplemental Figure 4). The *sepG^teton^* mutant, grown to mature hyphal growth under non-inducing conditions, appeared almost entirely aseptate indicating an essential role in septation (Supplemental Figure 4). To confirm a role for SepG in *A. fumigatus* hyphal septation, the *sepG^teton^* mutant was cultured to mature hyphal growth in the presence of increasing doxycycline concentrations to induce gene expression and then stained with calcofluor white to visualize septa. Under inducing conditions, septa were clearly visible throughout *sepGt^eton^*hyphae verifying SepG as a bona fide regulator of septation (Supplemental Figure 5).

Since our group and others have previously shown that even moderate defects in septation rate or septum morphology increase susceptibility to cell wall perturbing agents [2, 3, 33, 34], we next challenged the set of mutant strains with Congo Red (CR) or the echinocandin. caspofungin (Figure 6). Serial dilutions (10^6^-10^2^ conidia/ml) of each strain were spotted onto media containing 40 μg/ml CR and incubated for two days. Multiple mutant strains exhibited higher susceptibility to CR, including Δ*dopA*, Δ*cwf15*, and Δ*panA*, with *sepG^teton^*and Δ*pabA* unable to form colonies in this assay. To test caspofungin susceptibility, conidia were evenly spread on minimal media, a caspofungin drug strip was applied, and plates were incubated for two days. Compared to the inhibitory effect of caspofungin on the parental control (Δ*akuB-pyrg+*), Δ*cwf15*, Δ*rcoA*, *sepG^teton^* and Δ*pabA* appeared more strongly inhibited by caspofungin, with reduced growth in the zones of inhibition near the drug strip. Notably, *sepG^teton^* growth was completely inhibited in the presence of caspofungin concentrations above the minimum effective concentration, consistent with our previous results studying aseptate strains of *A. fumigatus* in this assay [2, 3].

**Figure 6:**
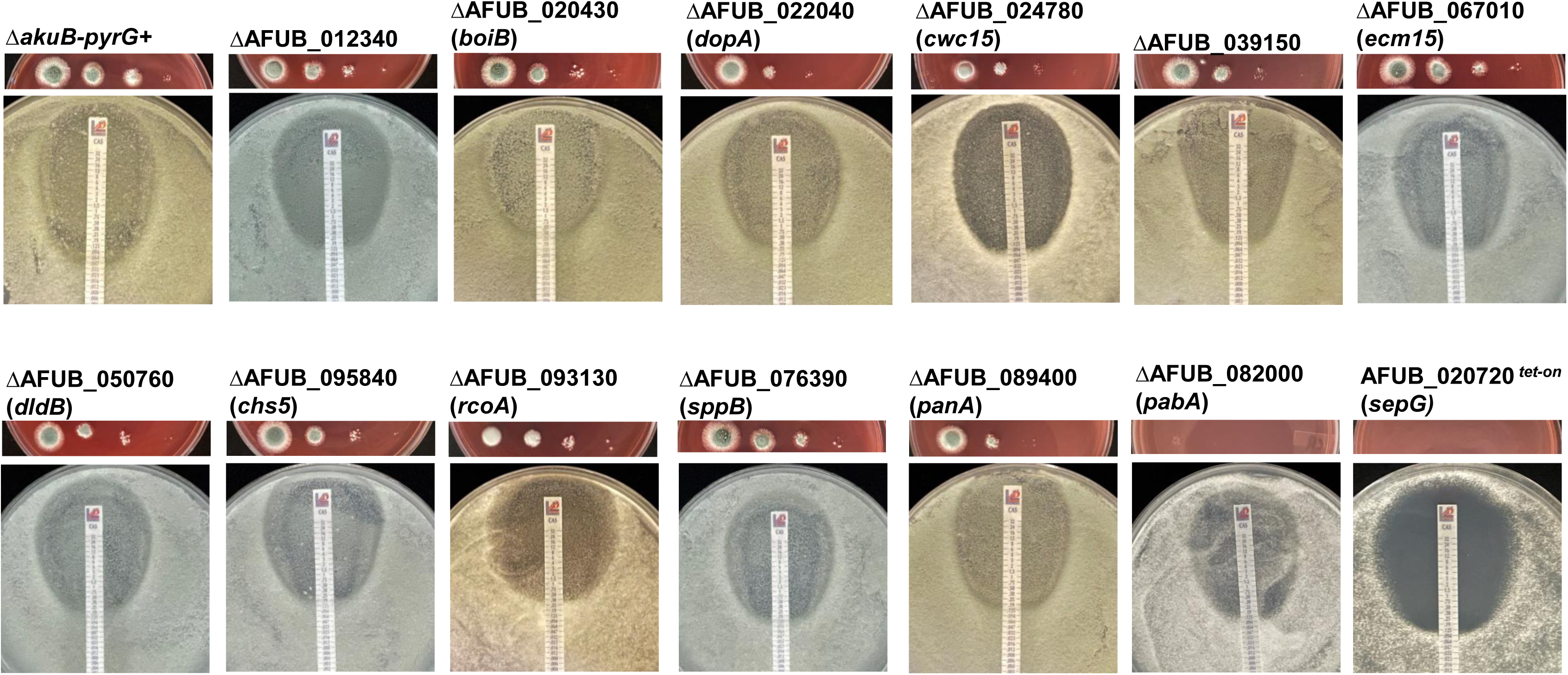
Cell wall stress characterization of selected mutants. Top: 10 μl of serial dilutions (10^6^ conidia/ml to 10^3^ conidia/ml) of strains were spot inoculated onto GMM plates supplemented with 40μg/ml congo red. Plates were incubated for 2 d at 37°C before imaging. Bottom: Drug strip diffusion assay using caspofungin drug strips. Strips were applied to plates after 10^6^ conidia of the indicated strain was spread evenly across a GMM plate. Plates were incubated for 48h at 37°C.

## Discussion

TurboID is increasingly used in fungi as a powerful tool to dissect pathways and cellular features involved in pathogenesis [16, 17]. Despite the limitations of this emerging technology, which are discussed below, we successfully utilized the terminal SIN components SidB and MobA to enrich for proteins with known roles in septation and identified proteins previously uncharacterized in *A. fumigatus* whose loss impacted septation and/or echinocandin susceptibility. Three notable examples are Cwc15, PabA, and SepG. Cwc15, previously uncharacterized in *A. fumigatus*, is predicted to be part of the Prp19 spliceosome. Orthologs of Cwc15 have been studied in *S. pombe, Saccharomyces cerevisiae* and the plant pathogen *Magnaporthe oryzae*, in which deletion mutants displayed defects in conidiation, cell wall integrity and virulence [31, 35]. Loss of *MoCWF15* resulted in a delay in appressorium formation, which has been linked to septation deficiency [36, 37]. We found loss of *cwc15* from *A. fumigatus* resulted in delayed septation, increased CR sensitivity and decreased growth under echinocandin stress by a drug strip assay. The *pabA* gene encodes a putative Protein Phosphatase 2A (PP2A) regulatory subunit. PP2A is a member of the <u>Stri</u>atin-associated <u>P</u>hosphatase <u>A</u>nd <u>K</u>inase complex (STRIPAK), which has been shown to regulate members of the SIN in both yeast and filamentous fungi [38, 39]. A different PP2A subunit, ParA, has been previously studied in *A. fumigatus* showing that *parA* deletion causes a radial growth defect, CR sensitivity and hyperseptation [40]. Similarly, the Δ*pabA* mutant is growth deficient, is inviable under CR stress and forms mature hyphae that appear hyperseptate. In *Sordaria macrospora*, STRIPAK was shown to be directly involved in modulating phosphorylation states of the SIN kinases [41]. ParA and PabA have each been previously characterized in *A. nidulans*, wherein a *pabA* mutant was severely growth restricted and displayed increased distance between septa, contrary to Δ*pabA* in the present study [29]. This suggests a diversion in function of PabA between the two species but emphasizes conserved necessity of STRIPAK for SIN regulation. Most striking among the genes studied here was the near complete loss of septation upon repression of *sepG* in the *sepG^TetOn^* strain. SepG is a conserved IQ-domain containing GTPase activating protein (IQGAP) which has been previously characterized in *A. nidulans* [32]. IQGAPs are known for their role in cytokinesis [42]. The *sepG* ortholog in *A. nidulans* was identified by a random mutagenesis screen to identify important players in cytokinesis [43]. Similar to our experience, a *sepG* deletion mutant in *A. nidulans* was not suitable for experimental manipulations. Instead, *sepG* was modulated using the *alcA* or *niaD* promoter. *A. nidulans sepG* suppression resulted in loss of septation and failure of contractile actin rings to constrict [32]. Using a doxycycline regulatable promoter system to repress *sepG* expression, we found consistent phenotypes, including a loss of hyphal septation corresponding to a loss of viability under caspofungin stress. Although it is unknown if any of the proteins studied here are direct effectors or regulators of the SidB/MobA complex, the septation-relevant phenotypes generated by their deletion suggest that TurboID analyses successfully identified SidB/MobA near-neighbor regulators of the septation process.

During our adaptation of TurboID to *A. fumigatus*, several limitations became evident. A primary factor we encountered is high background. This has been observed by other groups using streptavidin pulldowns, suggesting the background may be due to non-specific binding to magnetic beads or to streptavidin itself [44–46]. There are few proteins known to be biotinylated *in-vivo*; Svoboda et al. identified only 6 proteins carrying biotin moieties across all samples in their study using *A. nidulans*, including septin AspD [44]. Nonetheless, LC/MS-MS following streptavidin/streptactin pulldowns often identify hundreds to thousands of proteins in fungal lysates, regardless of how stringent blocking and elution procedures are [15, 17, 44, 45]. Western blots performed with crude extract and probed with streptavidin-HRP displayed bands that may correspond to known biotinylated proteins like pyruvate decarboxylase, but the presence of many bands here suggests the suite of natively biotinylated proteins in *A. fumigatus* may be quite extensive. Our streptavidin-HRP western blots confound those published previously using *A. fumigatus* mycelium [45]. Variations in extraction and gel loading conditions may contribute to this discrepancy, but baseline biotinylation is nonetheless considerable when designing a TurboID experiment. Notable endogenous biotinylation in CEA10 compelled us to introduce a control strain expressing cytoplasmic TurboID. In comparing our tagged strains to this control rather than a TurboID-free strain, we aimed to increase background noise such that the interaction partners of low-expression tagged proteins would be more significant in comparison. In light of this excess binding to streptavidin beads, further consideration should be given to treatment of samples after pulldown. Due to the strong interaction between streptavidin and biotin, (K_d_ ∼10^-14^) other groups have pre-blocked streptavidin beads or treated beads with harsh urea buffers after pulldown to remove proteins not engaged in this tight interaction [17, 44]. These steps seem to somewhat reduce background compared to our study.

Our study deviates from other TurboID studies in that previous studies rely on bait proteins with stable localizations, while our baits, SidB and MobA, migrate from the SPB to the site of septation [12, 16, 17, 44]. The translocation of TurboID-tagged bait proteins provides extra opportunity for TurboID to biotinylate proteins it may encounter during transport. We attribute the vast number of identified proteins in our pulldowns somewhat to this movement, though comparison to pOtef-TurboID rather than a TurboID-free strain may have reduced the impact of this non-specific biotinylation on our downstream analysis. Future work to identify critical septation regulators will involve using proteins with stable localizations either at the SPB or the site of septation to mitigate this potential source of noise. This source of noise may also be reduced by utilization of an inducible TurboID expression construct, such that TurboID-tagged bait is expressed for a shorter, more controlled duration than was performed here.

In the current study, we have adapted the TurboID system for proximity labeling to *A. fumigatus*. By overlapping datasets of two co-localizing proteins, we overcame the limitation of studying a protein without a stable cellular localization to identify novel septation effectors which may act upstream of the SIN at the spindle pole body or downstream at the septation site. This work helps clarify the local proteomes during septation and validates the use of TurboID to improve understanding of processes essential to drug susceptibility and virulence in fungal pathogens.

## Materials and Methods

### Strain generation and culture conditions

Strains used in this study are listed in Supplementary Table 1. Primers and CRISPR/Cas9 components used in this study are listed in Supplementary Table 2. Repair constructs for TurboID fusions were amplified from plasmid pAGRP-TurboID, made by cloning the coding sequence for TurboID into pAGRP using *BglII* and *SbfI* restriction enzymes [47]. Repair constructs for gene deletion were amplified from plasmid pJMR2 [48]. All mutants were made using a CRISPR-Cas9 gene editing system previously developed in our lab as described [49]. Transformation was performed using either CEA10 or its derivative Δ*akuB-pyrG+*, which is deficient for non-homologous end joining [50]. Proper integration of repair constructs was confirmed by diagnostic PCR, followed by Sanger sequencing for tagged strains. Unless otherwise listed, strains were cultured on Glucose Minimal Media and incubated at 37°C [51]. Conidia were harvested from 3-day-old cultures by flooding plates with sterile distilled water and scraping with a sterile cotton swab. Conidial suspensions were filtered through Miracloth, washed, and counted using a hemocytometer.

### Protein extraction and Western blots

10^8^ conidia were inoculated into 100 ml GMM broth supplemented with 50 μM biotin and incubated for 20 hrs at 37°C. Mycelium was filtered and washed with sterile distilled water, then flash-frozen in liquid nitrogen. Frozen mycelium was homogenized under liquid nitrogen using a mortar and pestle, and resulting powder was resuspended in lysis buffer supplemented with protease inhibitors (Pefabloc, Protease Inhibitor Cocktail) as described [23]. Protein quantification was determined by Bradford assay.

For western blots to visualize biotinylated proteins, mycelial powder was resuspended in RIPA buffer rather than the above lysis buffer. 50 μg of sample was denatured in Laemmli buffer + 5% β-mercaptoethanol for 5 min at 95°C. Denatured samples were allowed to cool before being run on a 7.5% protein gel (Biorad). Proteins were transferred to a PVDF membrane, blocked with 1% casein in TBS (Biorad), then incubated with Streptavidin-HRP (Sigma-Aldrich) diluted 1:20000 for 1 h.

For western blots requiring streptavidin pulldown, 2 mg total protein was incubated with 20 μl streptavidin magnetic beads for 1 h at room temperature. Proteins were eluted from the beads by boiling in Laemmli SDS buffer for 5 min at 95°C. 30 ul of eluted samples were run on a 7.5% protein gel, transferred to PVDF membrane, blocked with 1% casein in TBS, and blotted using an anti-TurboID primary antibody (Agrisera) or an anti-CotA specific antibody developed by our lab [23].

### Streptavidin bead precipitation and LC/MS-MS analysis

To isolate biotinylated proteins for LC/MS-MS analysis, crude protein lysates containing 3 mg total protein were incubated with streptavidin magnetic beads for 1 h at room temperature. Proteins were eluted off the beads in the presence of excess biotin. Samples were stored at-80°C until processed at the Duke University Proteomics and Metabolomics Core. For processing, samples were reduced for 15 min at 80°C, alkylated with 20 mM iodoacetamide for 30 min at room temperature, then supplemented with a final concentration of 1.2% phosphoric acid and 695 µL of S-Trap (Protifi) binding buffer (90% MeOH/100mM TEAB). Proteins were trapped on an S-Trap micro cartridge, digested using 20 ng/µL sequencing grade trypsin (Promega) for 1 hr at 47°C, and eluted using 50 mM TEAB, followed by 0.2% FA, and lastly using 50% ACN/0.2% FA. All samples were then lyophilized to dryness. Samples were resuspended in 12 µL of 1% TFA/2% acetonitrile with 12.5 fmol/µL of yeast ADH. A study pool QC (SPQC) was created by combining equal volumes of each sample. Quantitative LC/MS/MS was performed on 2 µL of each sample, using an MClass UPLC system (Waters Corp) coupled to a Thermo Orbitrap Fusion Lumos high resolution accurate mass tandem mass spectrometer (Thermo) equipped with a FAIMSPro device via a nanoelectrospray ionization source. Briefly, the sample was first trapped on a Symmetry C18 20 mm × 180 µm trapping column (5 μl/min at 99.9/0.1 v/v water/acetonitrile), after which the analytical separation was performed using a 1.8 µm Acquity HSS T3 C18 75 µm × 250 mm column (Waters Corp.) with a 90-min linear gradient of 5 to 30% acetonitrile with 0.1% formic acid at a flow rate of 400 nanoliters/minute (nL/min) with a column temperature of 55°C. Data collection on the Fusion Lumos mass spectrometer was performed for three difference compensation voltages (-40v,-60v,-80v). Within each CV, a datadependent acquisition (DDA) mode of acquisition with a r=120,000 (@ m/z 200) full MS scan from m/z 375 –1500 with a target AGC value of 4e5 ions was performed. MS/MS scans with HCD settings of 30% were acquired in the linear ion trap in “rapid” mode with a target AGC value of 1e4 and max fill time of 35 ms. The total cycle time for each CV was 0.66s, with total cycle times of 2 sec between like full MS scans. A 20s dynamic exclusion was employed to increase depth of coverage. The total analysis cycle time for each sample injection was approximately 2 hours. Data were imported into Proteome Discover 3.0 (Thermo Fisher) and peptides were aligned to the SwissProt *A. fumigatus* database. Raw mass spec data is available in ProteomeXchange under identifier PXD076255.

### Phenotypic characterization assays

To assess radial growth, 10 μl of a working water stock of 10^6^ conidia/ml was spot-inoculated into the center of a 100 mm petri dish with GMM agar. Data represent average growth of three transformants per strain. Plates were incubated at 37°C for 4 days. Colony diameter was measured every 24 hrs using calipers. To assess sensitivity to Congo Red, four serial dilutions (10^6^-10^2^ conidia/ml) of each strain were spotted onto media containing 40 μg/ml CR and incubated for 48 hrs at 37°C. Three transformants of each mutant and a control strain was included on each plate for direct comparison of mutant and parent. To measure susceptibility to echinocandin stress, 500 μl of sterile water containing 10^6^ conidia were evenly spread on a GMM agar plate and a caspofungin drug strip (Liofilchem) was applied after the plates dried. Plates were incubated for 48 hrs at 37°C. This assay was repeated with three transformants of each mutant.

### Fluorescence microscopy

To measure septation index, strains were inoculated glass-bottomed 24-well plates containing filtered GMM broth at a density of 10^4^ conidia/ml. Plates were incubated for 10 hours and samples were processed as previously described [3]. Briefly, samples were washed twice with PBS before being stained with 10 μg/ml calcofluor white in PBS (Fluorescence Brightener). After staining, samples were washed twice more with PBS before visualization using a Nikon TI2-A inverted microscope equipped with a Prime BSI express monochrome camera (Nikon, Tokyo, Japan). CFW staining was visualized using a DAPI filter.

## Acknowledgements

The authors thank Greg Wait, Tricia Ho and Erik Soderblom (Duke University Proteomics Core) for their expertise and assistance in LC-MS/MS data generation and analysis. This work was funded by NIH/NIAID awards R01 AI195581 and R01 AI158442 to J.R.F. and F31 AI191701 to H.T.

---

**Supplemental Figure 1:**
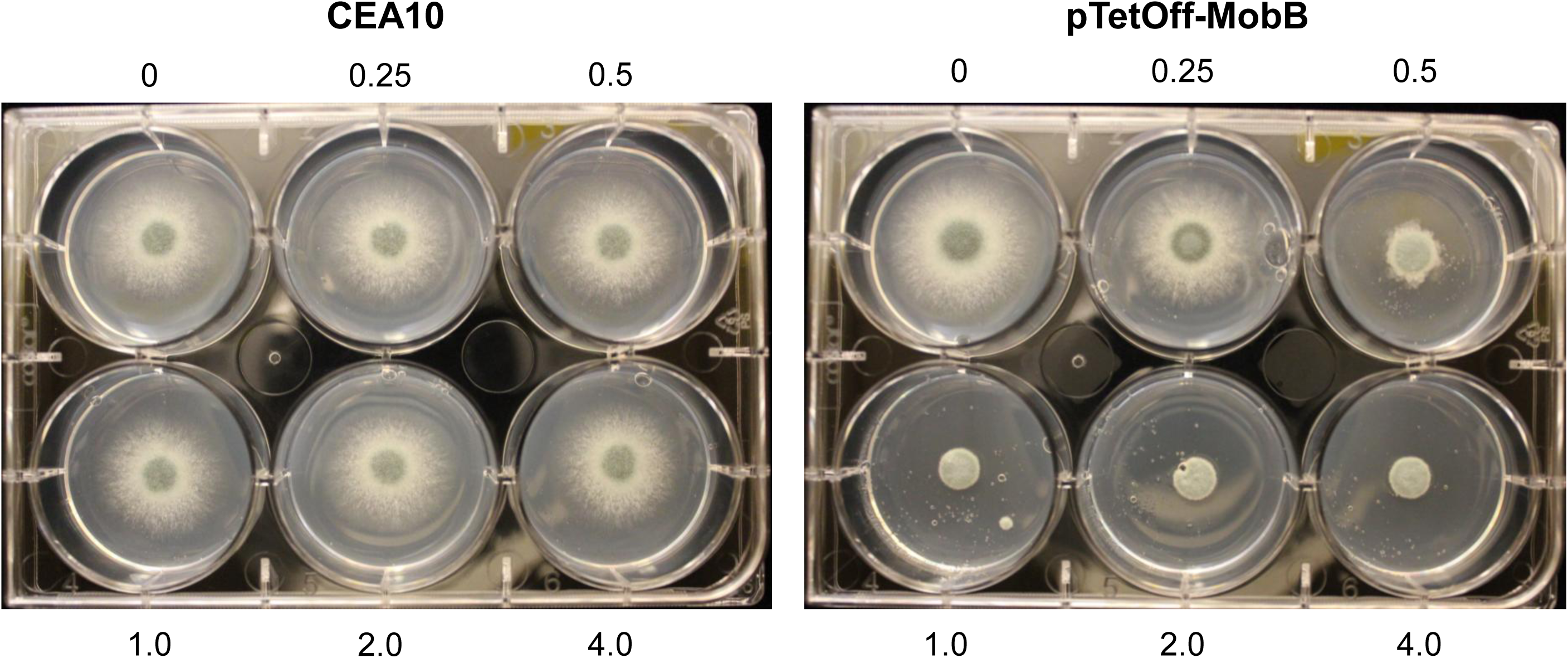
Impact of *mobB* repression on colony morphology. 10 μl of 10^6^ conidia/ml of the indicated strain was inoculated onto GMM in a 6-well plate containing increasing concentrations of doxycycline. Plates were incubated for 2 d at 37°C.

**Supplemental Figure 2:**
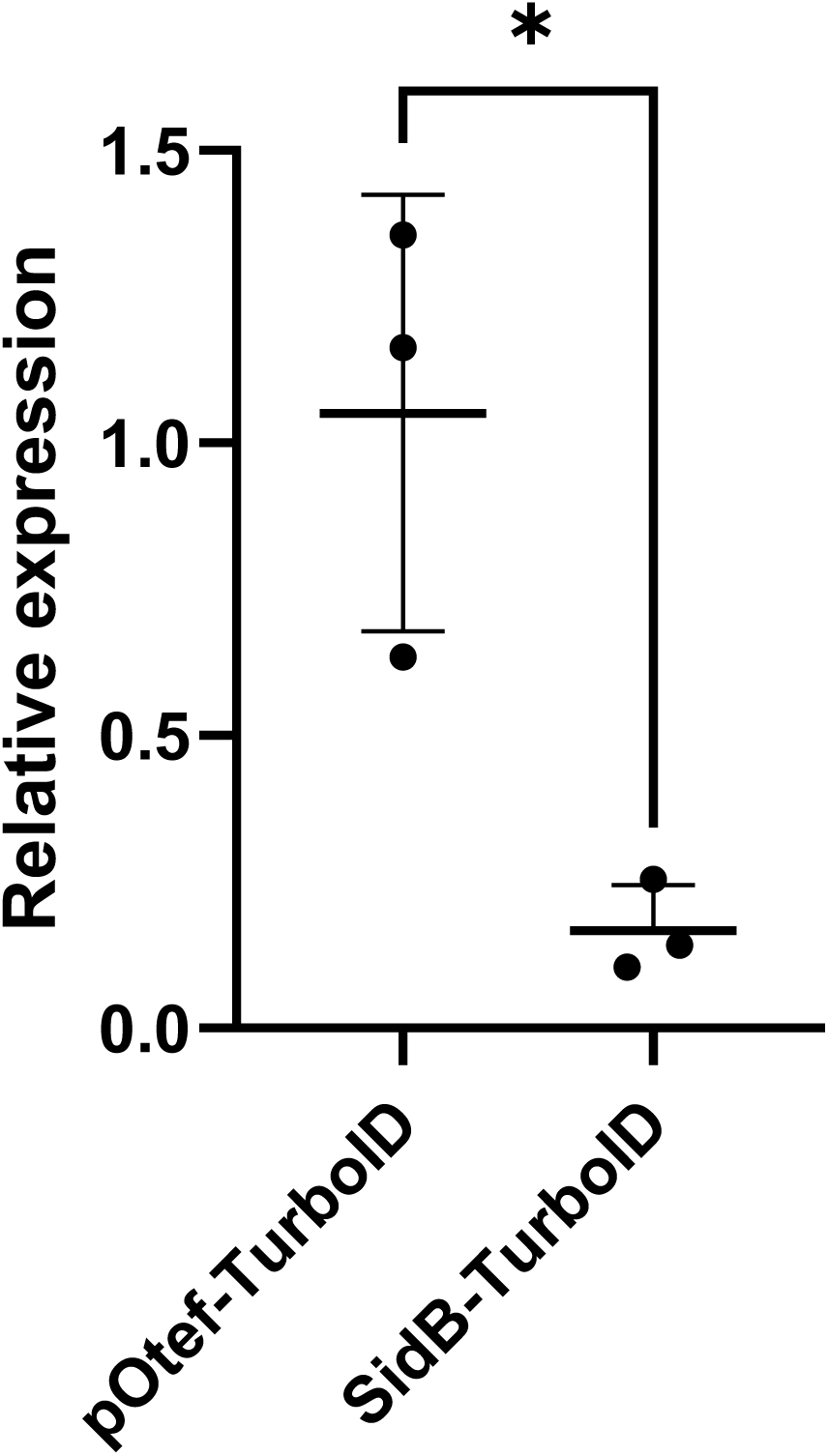
TurboID is highly expressed in pOtef-TurboID. RT-qPCR was performed using RNA extracted from overnight culture of the indicated strains. Expression was normalized to *tubA.* TurboID transcript level was compared to *pOtef-TurboID* by two-tailed T test. *: p<0.05. Data points are the average of two technical replicates of three biological replicates. C)

**Supplemental Figure 3:**
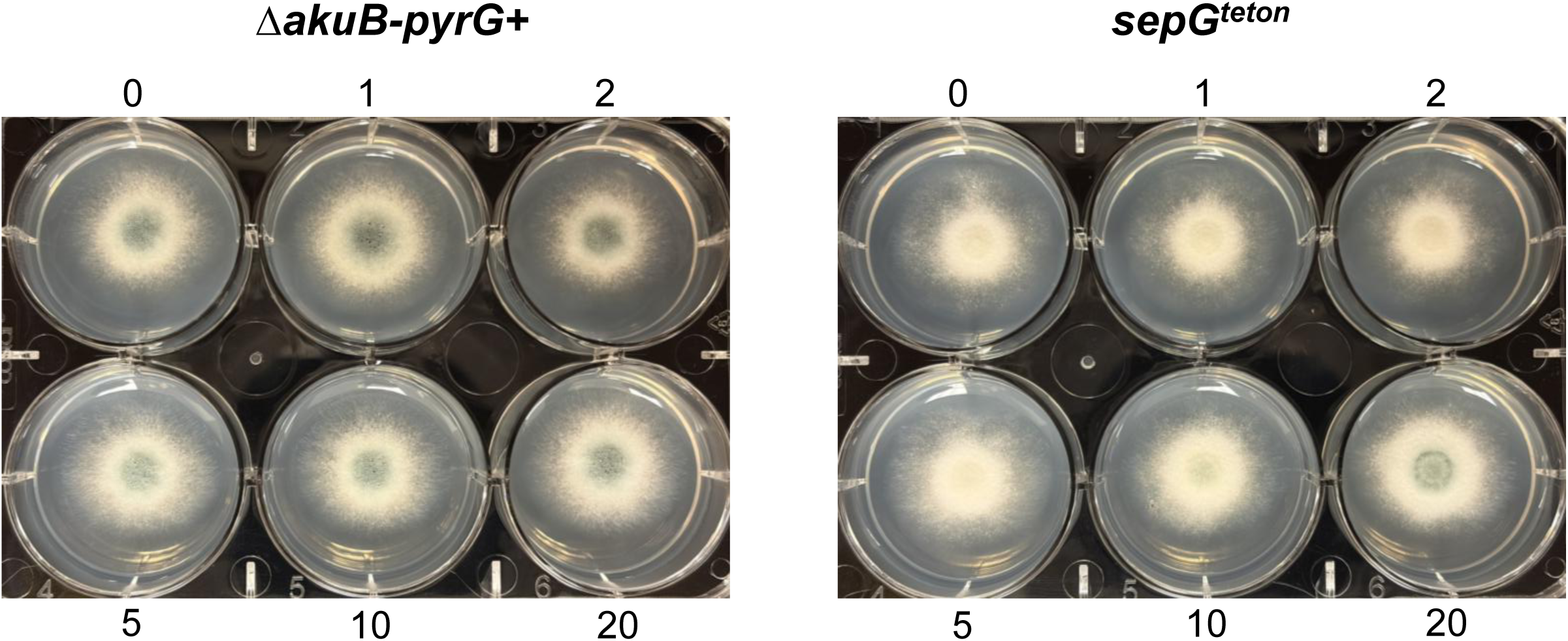
Impact of *sepG* repression on colony morphology and septation. A) 10 μl of 10^6^ conidia/ml of the indicated strain was inoculated onto GMM in a 6-well plate containing increasing concentrations of doxycycline. Plates were incubated for 2 d at 37°C.

**Supplemental Figure 4:**
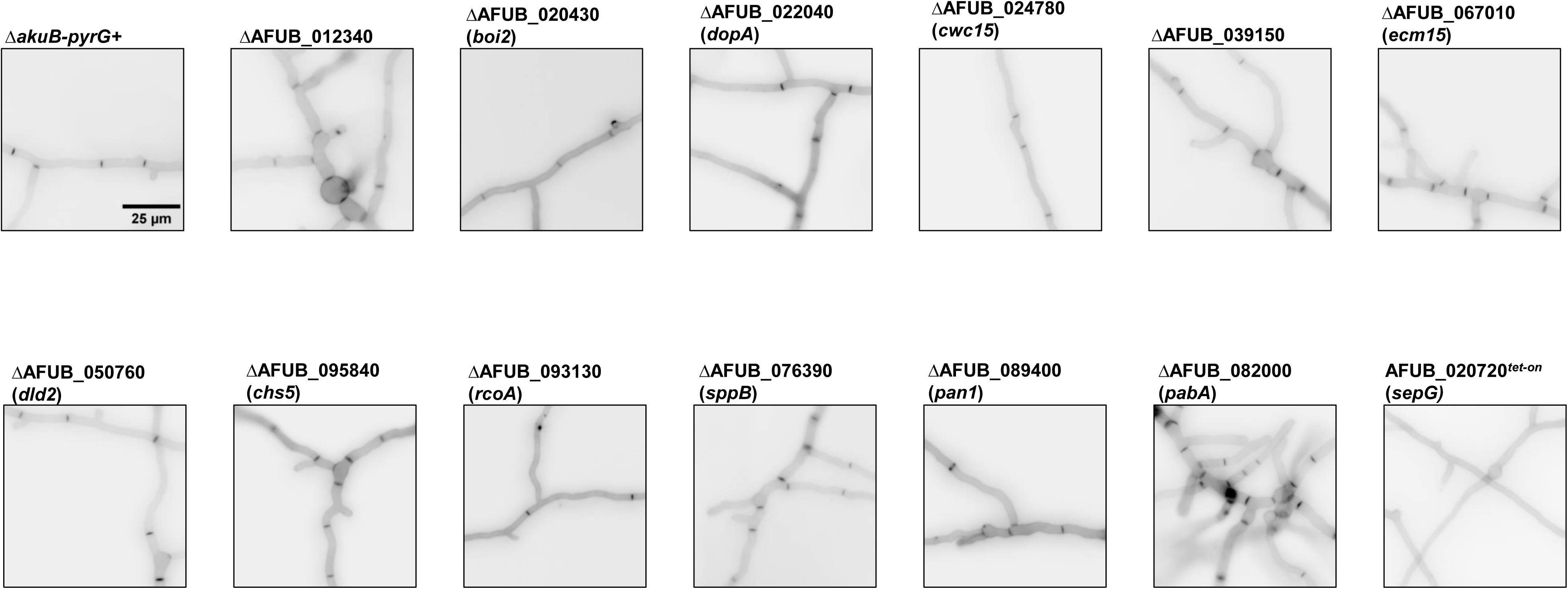
Hyphal septation of selected mutants. 10^4^ conidia of each strain was inoculated in 1ml GMM in a glass-bottomed 24 well plate and incubated for 16 h at 37°C. Wells were washed and stained with CFW to visualize septa.

**Supplemental Figure 5:**
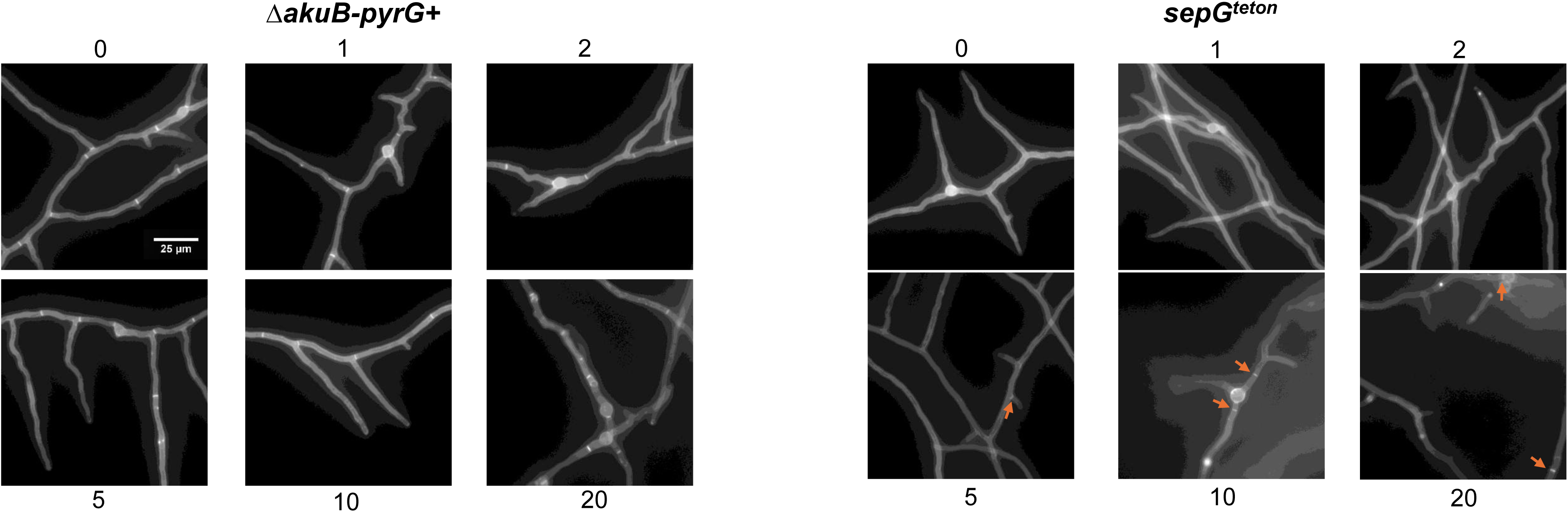
Induction of *sepG* expression restores septation. Conidia were inoculated into wells of a glass-bottomed 24-well plate in GMM supplemented with increasing amounts of doxycycline and incubated for 16 h at 37°C. Coverslips were stained with CFW and septa were imaged on an inverted fluorescent microscope using a DAPI filter.

